# Ultrafast Laser-Probing Spectrocopy for Studying Molecular Structure of Polymeric Proteins

**DOI:** 10.1101/081943

**Authors:** Huihun Jung, Chester J. Szwejkowski, Abdon Pena-Francesch, Benjamin Allen, Şahin Kaya Özdemir, Patrick Hopkins, Melik C. Demirel

**Affiliations:** Department of Engineering Science and Mechanics and Materials Research Institute, Pennsylvania State University, University Park, PA, 16802; Department of Mechanical and Aerospace Engineering, University of Virginia, Charlottesville, VA 22904; Huck Institutes of Life Sciences, Pennsylvania State University, University Park, PA, 16802; Department of Biochemistry and Molecular Biology, Pennsylvania State University, University Park, PA, 16802; Department of Electrical and Systems Engineering, Washington University, St. Louis, MO 63130

## Abstract

We report the development of a new technique to screen protein crystallinity quantitatively based on laser-probing spectroscopy with sub-picosecond resolution. First, we show theoretically that the temperature dependence of the refractive index of a polymeric protein is correlated to its crystallinity. Then, we performed time-domain thermo-transmission experiments on purified semi-crystalline proteins, both native and recombinant (i.e., silk and squid ring teeth), and also on intact *E. coli* cells bearing overexpressed recombinant protein. Our results demonstrate, for the first time, quantification of crystallinity in real time for polymeric proteins. Our approach can potentially be used for screening an ultra-large number of polymeric proteins *in vivo*.

## Introduction

Nature has optimized polymeric proteins through millions of years of evolution to provide diverse materials with complex structures (1). Thus, studying polymeric proteins at the molecular level will enable the efficient design of new materials that are finely tunable (2), flexible, biodegradable (3), self-healing (4) and have physical properties and functionalities different (5) or superior to those present in conventional materials (6). Polymeric proteins are characterized by long-range ordered molecular structures (*e.g.,* β-sheets, coiled coils, or triple helices) that arise due to highly repetitive primary amino acid sequences. These features promote the formation of a self-assembled structural hierarchy (7).

Polymeric (i.e., repetitive) proteins are key to the creation of many designer materials with programmable flexibility, biocompatibility, superior optical properties, energy efficiency, and mechanical strength. Characterization of polymeric proteins (or large macromolecular complexes) has been an interest not only for the molecular biologist to understand how biological macromolecules function, but also for the materials scientist to design novel materials and devices. Traditional techniques for probing protein structure include circular dichroism (CD), light scattering and fluorescence spectroscopy, Nuclear Magnetic Resonance (NMR), cryo-electron microscopy (cryo-EM), small-angle X-ray scattering (SAXS), and X-ray crystallography (8). However, there remains an urgent need for fast and efficient techniques that can screen the structural properties of large numbers of protein sequences with minimal sample volume or in living cells (9).

Here we report a new technique to characterize protein crystallinity based on ultrafast laser-probing spectroscopy. The time-domain thermo-transmission (TDTT) technique we describe here enables screening of polymeric proteins in milliseconds, a feat that would be impossible to achieve with existing approaches such as fluorescence, immunostaining, or functional assays (10). We studied three different semi-crystalline proteins and showed that crystallinity correlates quantitatively with transient thermo-optic response. Moreover, we demonstrated that the TDTT technique can work qualitatively *in vivo* by measuring the thermo-optical properties of overexpressed SRT protein in *E. coli.*

Silk (11) and SRT (12) are semi-crystalline polymeric proteins that form flexible, biodegradable, and thermally and structurally stable materials. The semi-crystalline morphology of these proteins, which originates from their β-sheet secondary structures, enables genetic tuning of their physical properties (13). Silk and SRT protein complexes are composed of several proteins of varying molecular weight (14). However this heterogeneity renders variable the physical properties of such samples. Therefore, we also studied a recombinant SRT (15) protein (Rec) with a unique molecular weight (i.e., 18 kDa) that forms a monodisperse material, providing advantages in physical properties (*e.g.,* melt viscosity, tensile strength, toughness).

## Results and Discussion

Native SRT (16) and silk (17) proteins were extracted from the tentacles of *Loligo vulgaris* and cocoons of *Bombyx mori* respectively following the protocol reported earlier. The proteins were dissolved in organic solvents and processed for deposition of pure polymeric protein films as described in Methods section. The recombinant 18 kDa SRT protein (15) was identified by a combination of RNA-sequencing, protein mass spectroscopy, and bioinformatics (i.e., transcriptome assembly), and then produced via recombinant expression in *E. coli*. Figure 1 shows optical and protein gel images for all three samples. Both silk and SRT provides a rich molecular architecture that can micro-phase separate to form amorphous Gly-rich and crystalline β-sheet regions with crystalline domain dimensions of 2-3 nm. However, SRT proteins show considerable diversity (variable AVSTH-rich) in their crystal-forming sequences (7), while silk protein is made up of highly ordered β-sheet stacks containing oligomers of GAGAGS repeats (18). To provide crystallinity values for comparison, we analyzed samples of Rec, SRT and silk protein by X-ray diffraction (XRD). The cast silk sample exhibited the lowest crystallinity, while the native SRT protein displayed lower crystallinity compared to Rec protein at room temperature (Fig. S1). The Rec, SRT and silk proteins were also analyzed using infrared spectroscopy (FTIR). The amide regions of the FTIR spectra confirm that the crystalline regions of all three proteins are composed of β-sheets and α-turns (Fig. S2).

**Figure 1.**
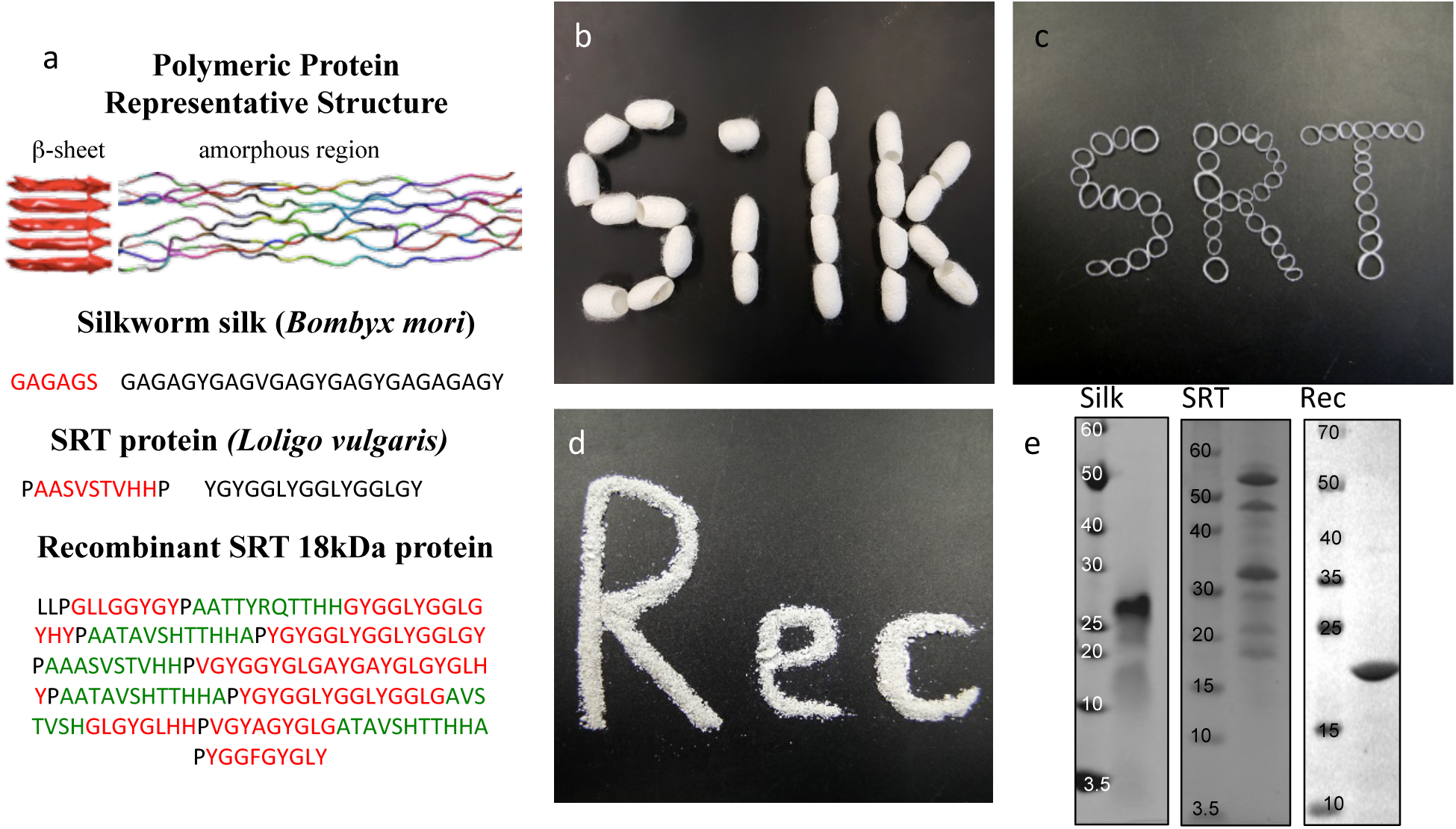
Polymeric Protein Characterization. (a) The assembly of β-sheet structures forms crystalline domains in repetitive silk and SRT proteins. Amorphous repetitive domains are stretched in the silk structures, but random in SRT. Segmented copolymer architecture of the recombinant 18kDa SRT protein sequence is marked as red (Gly-rich), green (AVSHT-rich), which are separated by proline residues (bold). (b)-(d) Optical images of silk cacoons, SRT rings, recombinant SRT (Rec) powder proteins. (d) SDS-Page shows protein molecular weight for degummed silk (varying between 10-30 kDa), for native SRT (varying between 15-55 kDA), and for recombinant SRT (18 kDa + 2 kDa histidine-tag).

The temperature dependence of the refractive indices (i.e., thermo-optic coefficient) of the polymeric proteins was reported earlier based on optical cavity measurements (19). In general, the thermo-optic coefficient can be written as a function of density change and temperature change, *dn*/*dT* = (*∂n*/*∂ρ*)_*T*_ (*∂ρ*/*∂T*) + (*∂n*/*∂T*)_ρ_ = −(*ρ∂n*/*∂ρ*)_T_ *α* + (*∂n*/*∂T*)_ρ_, where *ρ* is the density, *α* is the coefficient of thermal expansion. The first term on the right side of the equation is the volume-dependent thermal response, which is related to thermal expansion. The second term, on the other hand, is the volume independent thermal response that is solely determined by the electronic structure of the material. We measured the thermal expansion coefficient, *α*, for all protein samples (−95×10^−6^ ± 7×10^−6^ K^−1^ for SRT proteins and −300×10^−6^ ± 10×10^−6^ K^−1^ for silk). Apparent negative thermal expansion coefficients are not surprising for polymeric proteins. For rubbery materials, the thermal expansion coefficient depends on the strain. In general, the total expansion is obtained by adding the expansion coefficient, α, to the unstressed material, *α*’ = *α* - *ε/T*. For example, a natural rubber stretched three times of its length gives a contribution of about −10^−2^ K^−1^ at room temperature (20). Similarly, silk shrinks significantly in size when hydrated due to inter-domain stresses (*i.e.,* super-contraction of amorphous domains) in the native form (21).

For conventional polymers, the volume independent index change, (*∂n*/*∂T*)_ρ_, is small enough to be ignored. Hence, Zhang *et al.* (22) showed a linear relationship between *dn/dT* and *α* values by ignoring the volume independent term for conventional polymers. However, volume-independent refractive-index changes for polymeric proteins must not be ignored since crystalline β-sheet domains act as grain boundaries and significantly contribute to optical properties including reflectivity. For polymeric proteins, we show below that the volume-independent index-change term has negative sign (*∂n*/*∂T* < 0), leading to a stronger negative thermo-optic response.

We measured the relative magnitude and sign of the volume-independent thermo-optic coefficient, (*∂n*/*∂T*)_ρ_, using time-domain thermo-transmission (TDTT) (Fig. 2a). TDTT is centered upon concepts of ultrafast laser pump-probe spectroscopy, where, in TDTT, the measured transmission signal is related to the temporal decay of the probe pulse due to the transient energy density and transport mechanisms induced from the modulated pump pulse. The pump is used to trigger a rapid thermal process in the sample and the probe beam is used to examine the excited relaxation dynamics and energy changes of the excited volume. The temporal evolution of this process is monitored by varying the relative delay time between the pump and probe pulses. While the rapid absorption of the pump pulse leads to both temperature changes and thermal expansion, these signatures of the optical changes of the sample are separable in the time domain (23). The characteristic signatures of the temporal thermal response are a rapid exponential decay in ~2 ps after laser absorption, representing nonequilibrium redistribution and coupling among thermal carriers, followed by a slower exponential decay, representing the change in the absorption of the sample that is linearly related to temperature (i.e., diffusive thermal transport) (24).

**Figure 2.**
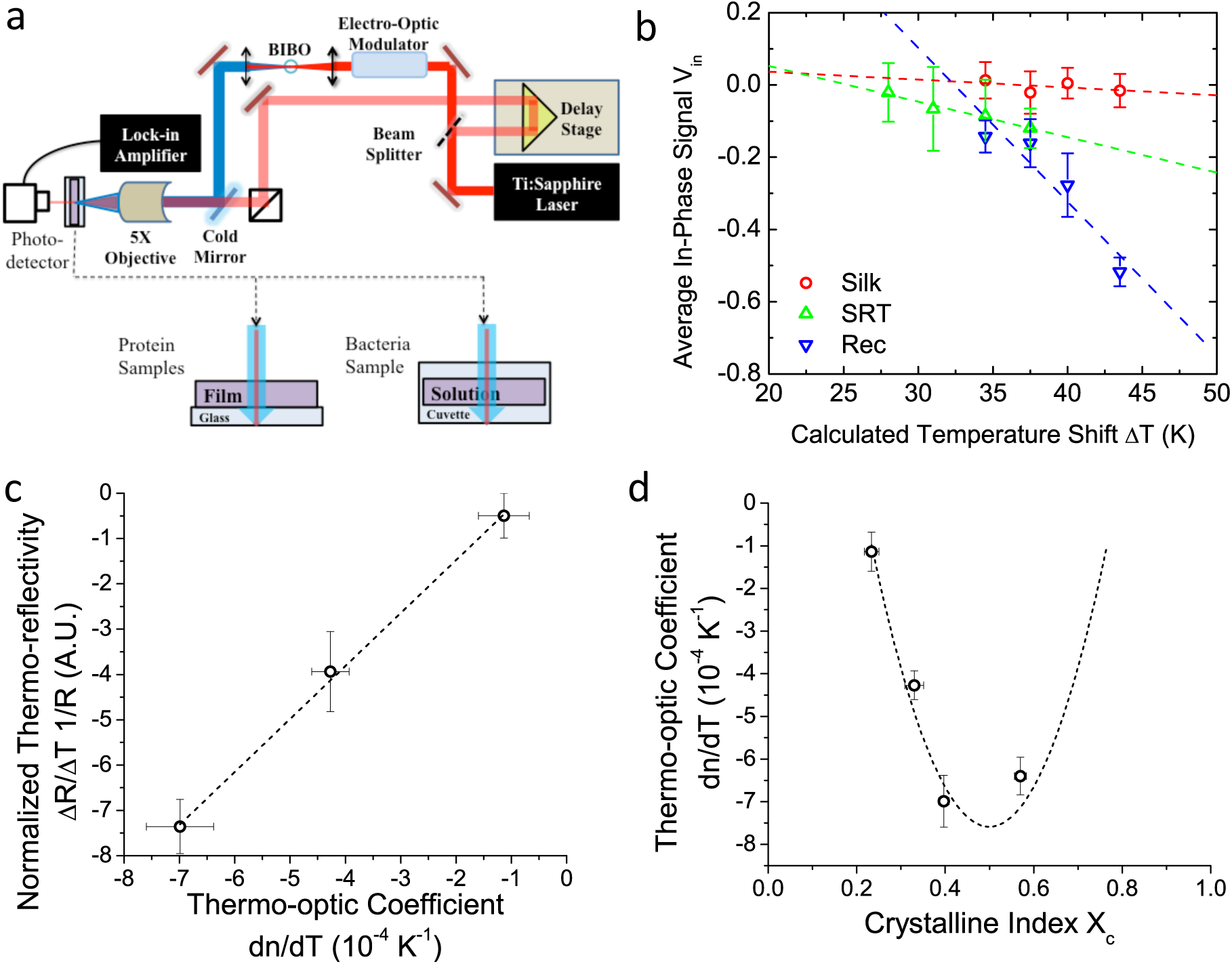
Transient thermo-optic set up and protein measurements. (a) Schematic of typical TDTT set up. The measured change in reflectivity is representative of the change in refractive index due to a change in temperature (dn/dT). (b) Average in phase signal (V_in_) due to a heating event from a pump laser pulse as a function of the temperature is representative of the change in reflectivity (R = 1 - Tr) due to a change in temperature. (c) The normalized thermo-reflectivity increases linearly with the thermo-optic coefficient. The thermo-reflectivity is calculated as (1/R) (dR/dT) = (√*2*/GQ)[V_∂_/(∆TV_0_)], where G is the gain of the preamplifier, Q is the quality factor of the circuit, Vo is the average dc voltage generated by the photodiode detector from pump-probe experiments. The slope of the dashed line is ~1.2, which agrees well with the theoretical prediction of 1.4 (See Supplementary Information) (d) Effect of the crystallinity index on the thermo-optic coefficient measured by all-protein WGM resonators. Methanol treated SRT has increased crystallinity (e.g., X_c_ increased from 0.34 to 0.58). The dashed line shows theoretical prediction for thermo-optic coefficient (e.g., dn/dT~[X_c_ (1 − X_c_)]) as a function of crystallinity.

Our measurements are performed outside the picosecond interval, which also ensures that our measurements avoid any non-linear optical effects from simultaneous pump and probe absorption in the sample. Using transmission geometry to detect the transient absorption changes (25), our measurements on three different polymeric proteins are then related to the volume independent term of the thermo-optic coefficient, as described in Supplementary Information (Fig. S3a-c). We benchmark our measurements and analysis using a thin gold film (Fig. S3d), as the thermo-optic coefficients of gold films have been extensively characterized (26).

First, monitoring the volume-independent reflectivity signal, V_in_, as a function of pump-laser power shows linear relationships with distinct slopes for each of the different protein materials (Fig 2b). These data are used to compute the thermo-reflectivity of each sample, which correlate linearly with the thermo-optic coefficients of the same materials (Fig. 2c) that we measured in our previously published work (19). Plotting the thermo-optic coefficients calculated from the experimental data for the tested proteins as a function of crystallinity revealed that the thermo-optic coefficient becomes more negative with increasing crystallinity index of the protein (Fig. 2d). Considering the Fresnel and Beer laws (see derivation in Appendix), an analytical relationship can be derived *dn/dT* = [*a z*(*n* + 1)^3 ∝/2(n − 1)] *X*_*c*_ (1 − *X*_*c*_), where *a* is the extinction coefficient, z is the film thickness, and *X*_*c*_ is the crystallinity index (Fig 2d), which agrees well with the experimental data.

In Figure 3, we demonstrate that the TDTT technique could work qualitatively *in vivo* by measuring the relative temperature changes in the thermo-optical properties of overexpressed recombinant protein SRT protein in *E. coli.* FTIR analysis of the overexpressed *E. coli* (Fig. S4a) reveals higher β-sheet content (48%) compared to purified recombinant 18kDa protein (38%). Typical total protein concentration in *E.coli* is estimated as 190 mg/ml, which corresponds to dry weight of 55% or volumetric composition of 40% (27). Hence, larger β-sheet content in *E. coli* compared to purified recombinant film is expected. On the other hand, as a negative control, we also performed FTIR studies (Fig. S4b) on the same empty vector strain of *E. coli*, which shows 6% lower β-sheet content (42%) compared to overexpressed *E. coli*. We attribute this difference to overexpressed SRT protein, which should contribute 6-8% increase to β-sheet content given that overexpression yields 15-20% excess recombinant SRT protein in *E. coli*. These results are also consistent with the TDTT results. Although the reflectivity data in bacterial measurements cannot be converted into exact crystallinity values (due to lack of thermo-optic coefficient for *E. coli*), it is clear that the reflectivity response correlates linearly with the laser power (~ temperature shift).

**Figure 3.**
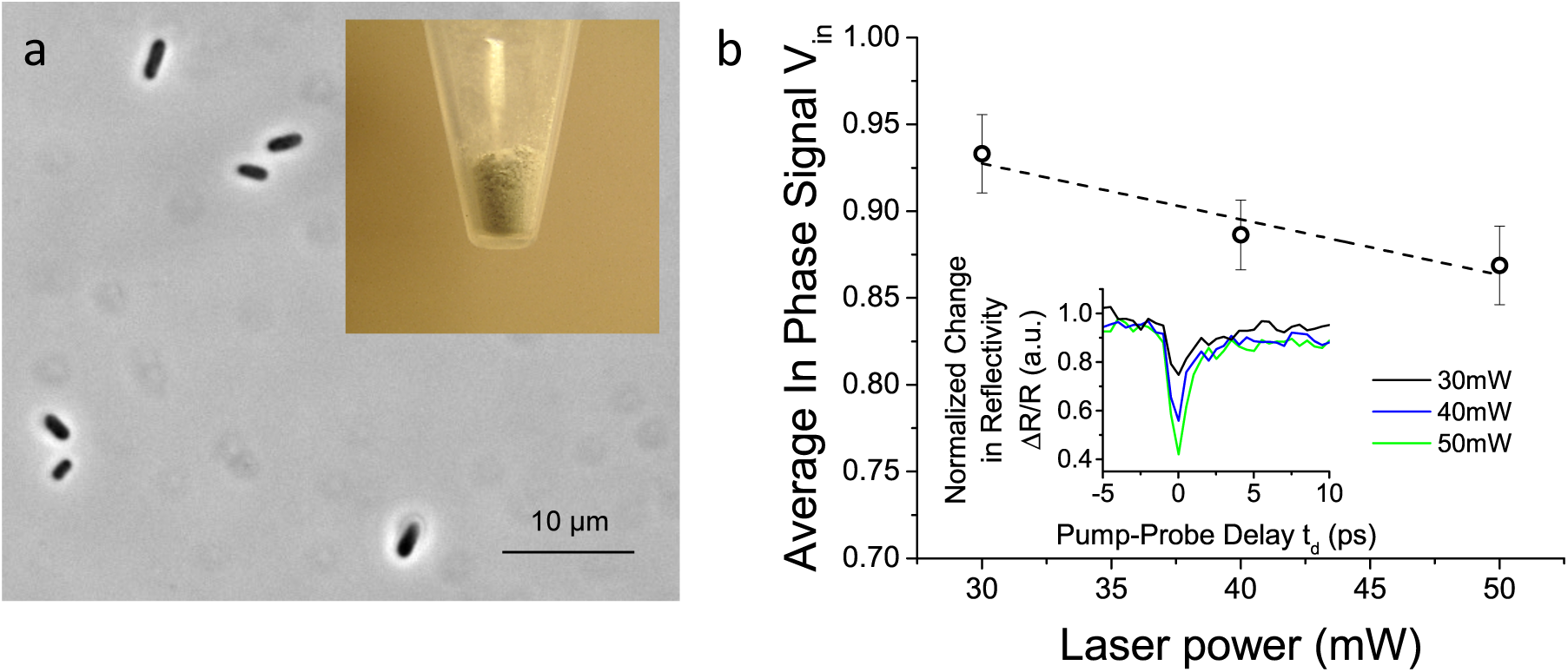
Bacterial TDTT Measurements. (a)Intact E. coli cells (strain BL21-DE3) that have recombinant SRT proteins (inset shows freeze-dried bacterial cells) (b) Normalized change in reflectivity for overexpressed recombinant protein SRT protein in E. coli as a function of laser power.

## Conclusion

We report the development of a novel optical spectroscopy technique to characterize the nanoscale morphology of polymeric proteins. We also show theoretically that the reflectivity measurement is correlated to the protein crystallinity via the thermo-optic coefficient. As we show here, time-resolved changes in the refractive index of semicrystalline proteins vary as a function of temperature, and the strength of this effect correlates with crystallinity. Ultimately, this allows us to quantify rapidly the crystallinity of a protein sample using time-domain thermo-transmission spectroscopy by decoupling volumetric thermal expansion from its structural response at room temperature. Our approach can potentially be used for screening an ultra-large number of polymeric proteins *in vivo*. The size of the library for these proteins is simply limited by the fluidic and electronic components of the sorting since the TDTT technique operates in the order of picoseconds. If this screening technique is achieved, we could answer many fundamental questions in polymeric protein research, such as (i) what is the underlying sequence-structure relationship for polymeric proteins? (ii) what is the complete set of semi-crystalline proteins, given that only few (*e.g.,* SRT and silk) have been discovered up to date. Successful development of this technique for polymeric proteins will have a significant impact on multiple applications in various fields (*e.g.,* materials science, agriculture, neurological diseases and medicine) and open new avenues of protein research.

## Materials and Methods

### Native squid ring teeth (SRT) and Silk

European common squid (*Loligo vulgaris*) were caught from the coast of Tarragona (Spain). The squid ring teeth (SRT) were removed from the tentacles, immediately soaked in deionized water and ethanol mixture (70:30 ratio v/v) and vacuum dried in a desiccator. Silk cocoons were purchased from Amazon, Inc, and we followed the protocol reported earlier (6).

### Expression and characterization of recombinant squid ring teeth (SRT) proteins

The SRT protein family is comprised of SRTs of different size that exhibit different physical properties. Heterologous expression of the smallest (~18 kDa) SRT protein extracted from *Loligo vulgaris* (LvSRT) was performed using the protocols described earlier (28). Briefly, the full length sequence was cloned into Novagen’s pET14b vector system and transformed into *E. coli* strain BL21(DE3). Recombinant SRT expression was achieved with a purity of ~90% and an estimated yield of ~50 mg/L. The yield increases approximately ten fold (i.e., ~0.5 g/L) in auto induction media. The size of the protein was confirmed via an SDS-page gel.

### Protein casting

SRT solution (either recombinant or native protein) was prepared by dissolving 50 mg/mL of protein in hexafluoro-2-propanol (HFIP). The solution was sonicated for 1 hour and vacuum-filtered in a 4-8 µm mesh size filter. 80 µL of SRT solution were poured into a PDMS toroid mold in successive 20 µL additions 1 minute apart. After the last addition, the HFIP was evaporated at room temperature in the fume hood for 5 minutes and the mold-SRT system was immersed in butadiene (plasticizer) at 80°C (above T_g_) for 30 minutes. The thermoplastic SRT film was peeled off with tweezers and excess butanediol was removed by rinsing with ethanol. For silk protein solution, we followed the protocol reported earlier (6). Aqueous silk solution was injected into the PDMS mold and evaporated at room temperature in a desiccator overnight.

### Pump-probe transient absorption measurements

A Ti:sapphire laser (Spectra-Physics, Tsunami) is used for time domain transmissivity measurements. Laser characteristics were 800 nm central wavelength, 10 nm bandwidth, 90 fs pulse duration, and 80 MHz reputation rate. The beam is split into two paths (pump and probe paths) by a polarizing beam splitter (PBS) and the pump is frequency doubled to 400 nm wavelength, pump: 7.5 mW (average power of pump modulated at 10 MHz using a sinusoidal envelope), probe: 2.5 mW of average power. The beam size 17 micron for the probe and 35 micron for the pump with synchronous sampling via mechanical delay stage. The transmittance signal is digitalized using a Thorlabs DET10A Si Detector and a Zurich Instruments UHFLI digital lock-in amplifier. The resulting voltage is on the order of 10^−6^ V; data was collected up to pump-probe time delays of 5.5 ns, with variation and noise of 1 nV and 0.2 µV respectively. The bandwidth of the detector was 1 GHz. The delay between pump and probe is mechanically controlled using Newport Optical Delay Line Kit. A combination of LabView and MatLab programs (developed in-house at the ExSiTE Lab, UVa) were used to analyze the data.

## Acknowledgments

MCD, BDA, AP, and HJ were supported by the Army Research Office under grant No. W911NF-16-1-0019, and Materials Research Institute of the Pennsylvania State University. PEH and CJS were supported by the Air Force Office of Scientific Research under grant No. FA9550-15-1-0079 and Army Research Office under grant No. 151630-101-GG11945-31345. We thank Mr. Huzeyfe Yilmaz for providing thermo-optic coefficient for polymeric proteins studied in this manuscript.

## Author contributions

MCD conceived the idea and derived structure-property equations. MCD, SKO, PH and BDA planned and supervised the research. AP fabricated the protein films, CS performed the pump-probe transient absorption measurements, and HJ worked on the cloning, recombinant expression and purification of polymeric proteins. All authors contributed to writing and revising the manuscript, and agreed on the final content of the manuscript.

## Additional information

The authors declare no competing financial interests. Correspondence and requests for materials should be addressed to MCD.

## SUPPLEMENTARY INFORMATION

**Figure S1.**
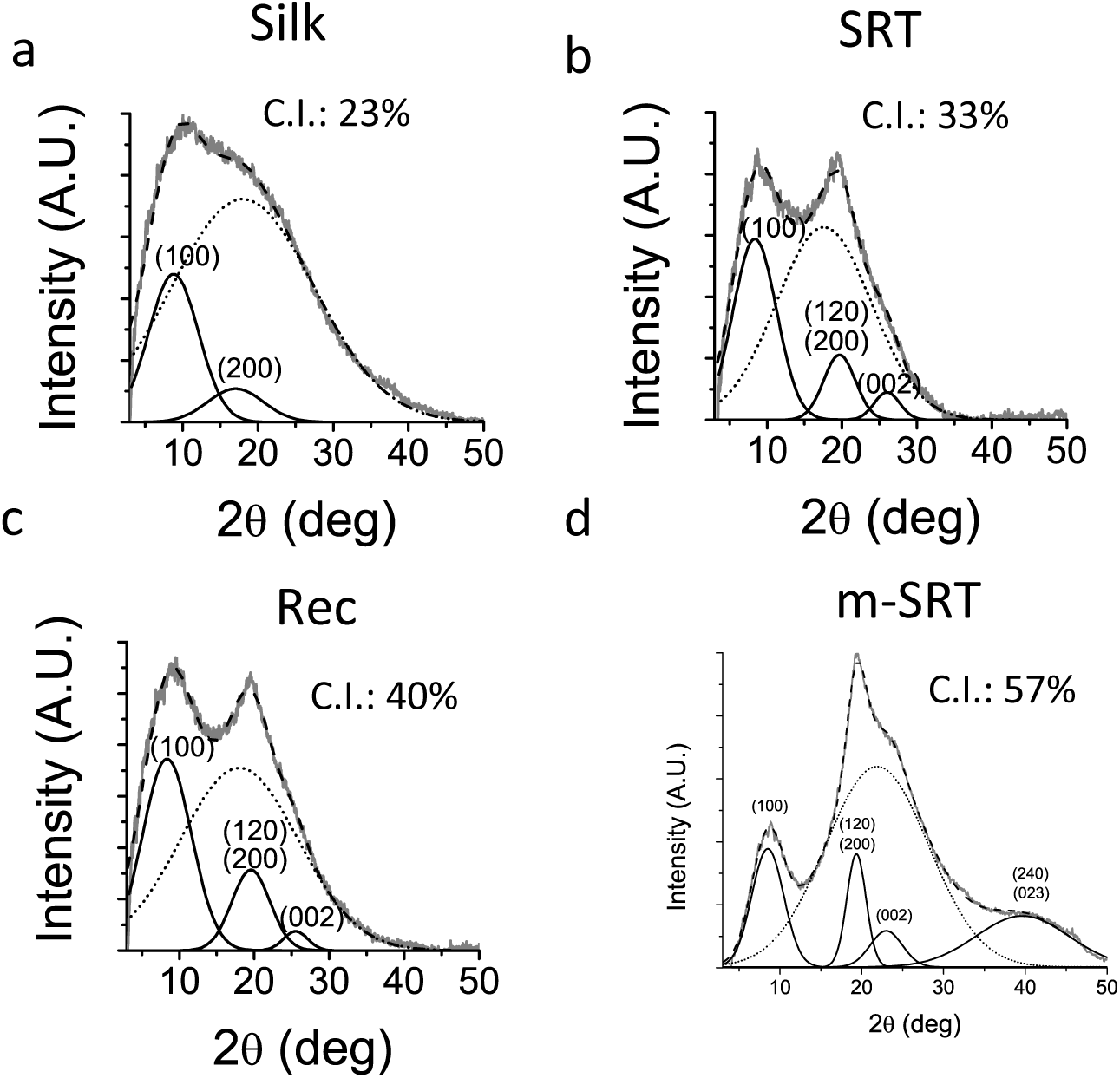
XRD (a, b, c, d) spectra for silk, native SRT, recombinant SRT (Rec) and methanol treated SRT (m-SRT) proteins show a semicrystalline profile where crystalline peaks (solid line) and an amorphous halo (dotted line) are fit.

**Figure S2.**
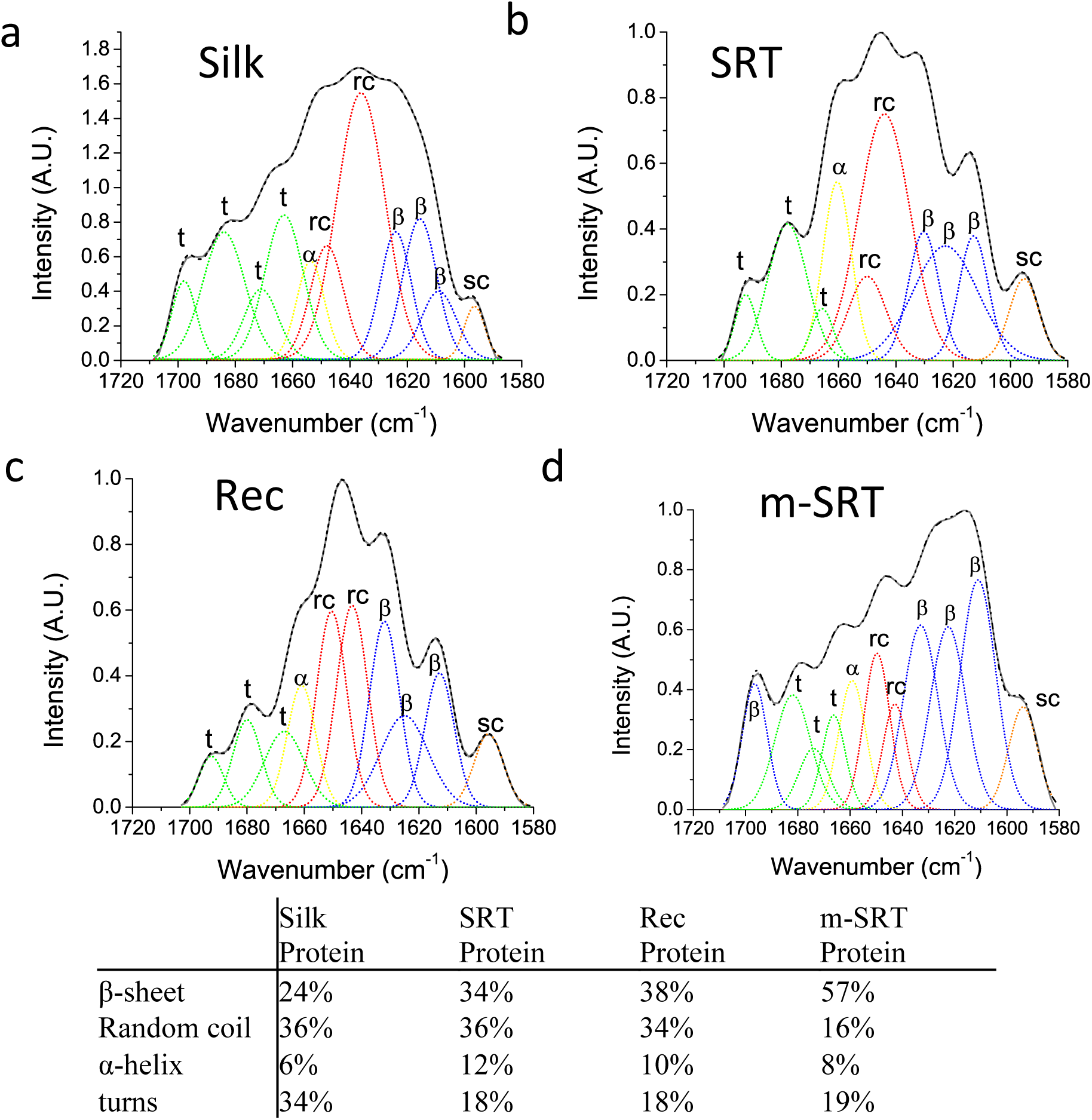
Deconvoluted FTIR (a, b, c, d) spectra for silk, SRT, Rec, and m-SRT are shown. The Amide-I band (1600-1700 cm^−1^) is representative of the secondary structures of the respective proteins. Based on the peak area, contribution of secondary structures is estimated in the table. Crystallinity of proteins in FTIR spectra is estimated from β-sheet content.

**Figure S3.**
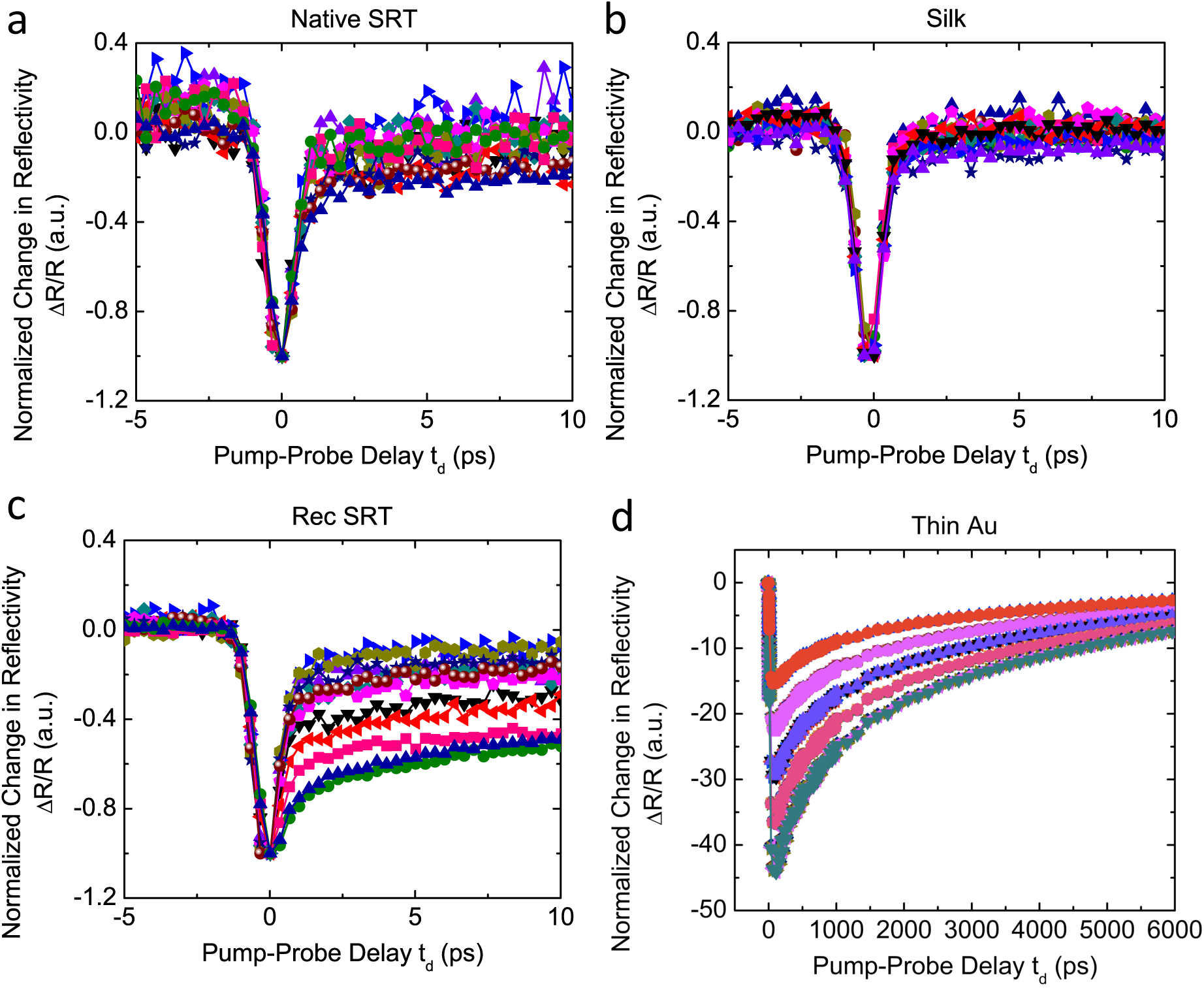
Normalized change in reflectivity due to a heating event from a pump laser pulse as a function of the delay time between the pump and probe beams for native SRT (a), silk (b) and recombinant SRT (c) proteins. The measured change in reflectivity is representative of the change in refractive index due to a change in temperature (dn/dT). (d) Control experiment with Au thin film.

**Figure S4.**
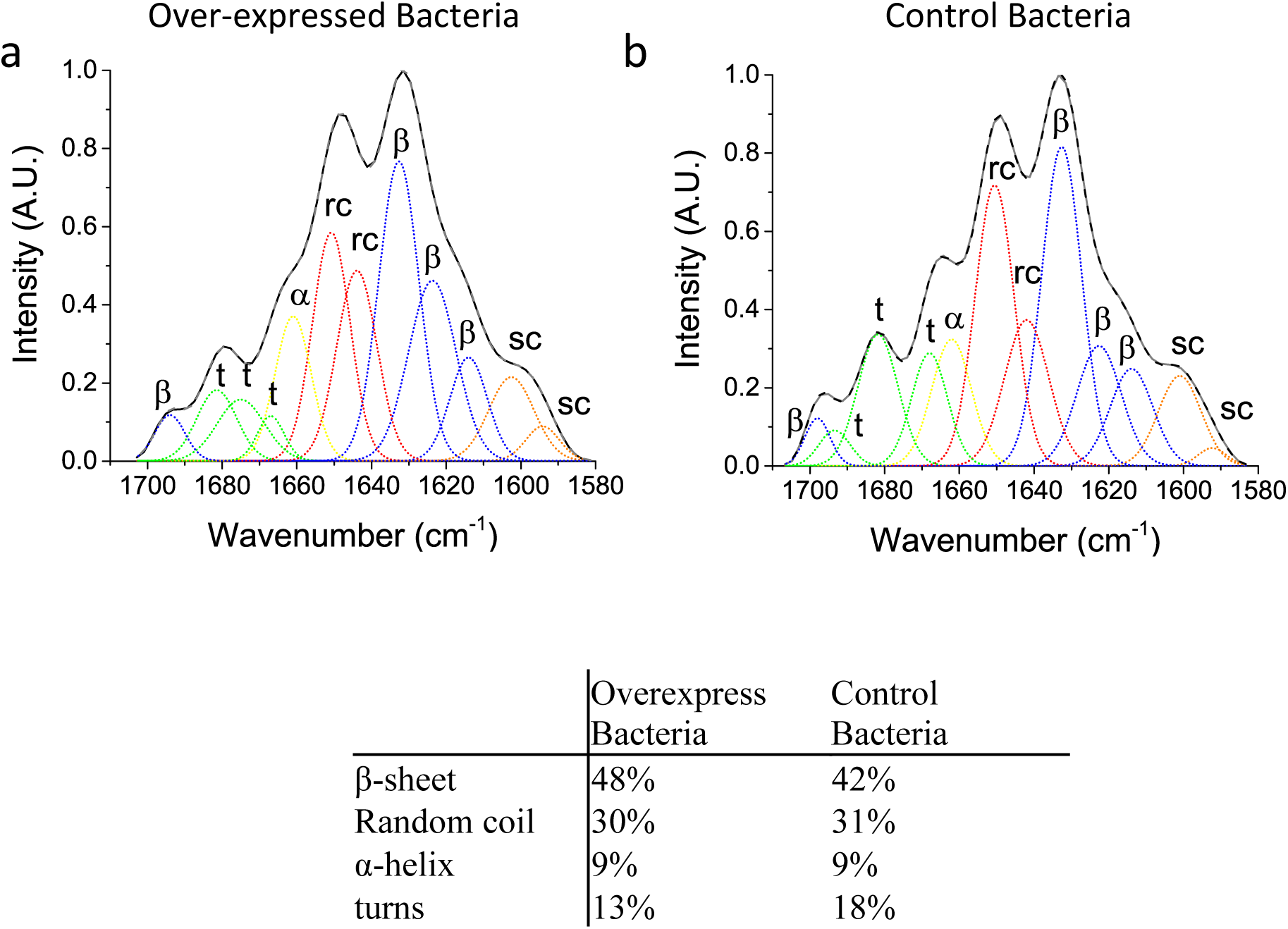
Deconvoluted FTIR (a, b) spectra for overexpressed and empty vector (control) *E. Coli* are shown. The Amide-I band (1600-1700 cm^−1^) is representative of the secondary structures of the respective proteins. Based on the peak area, contribution of secondary structures is estimated in the table.

**Figure S5.**
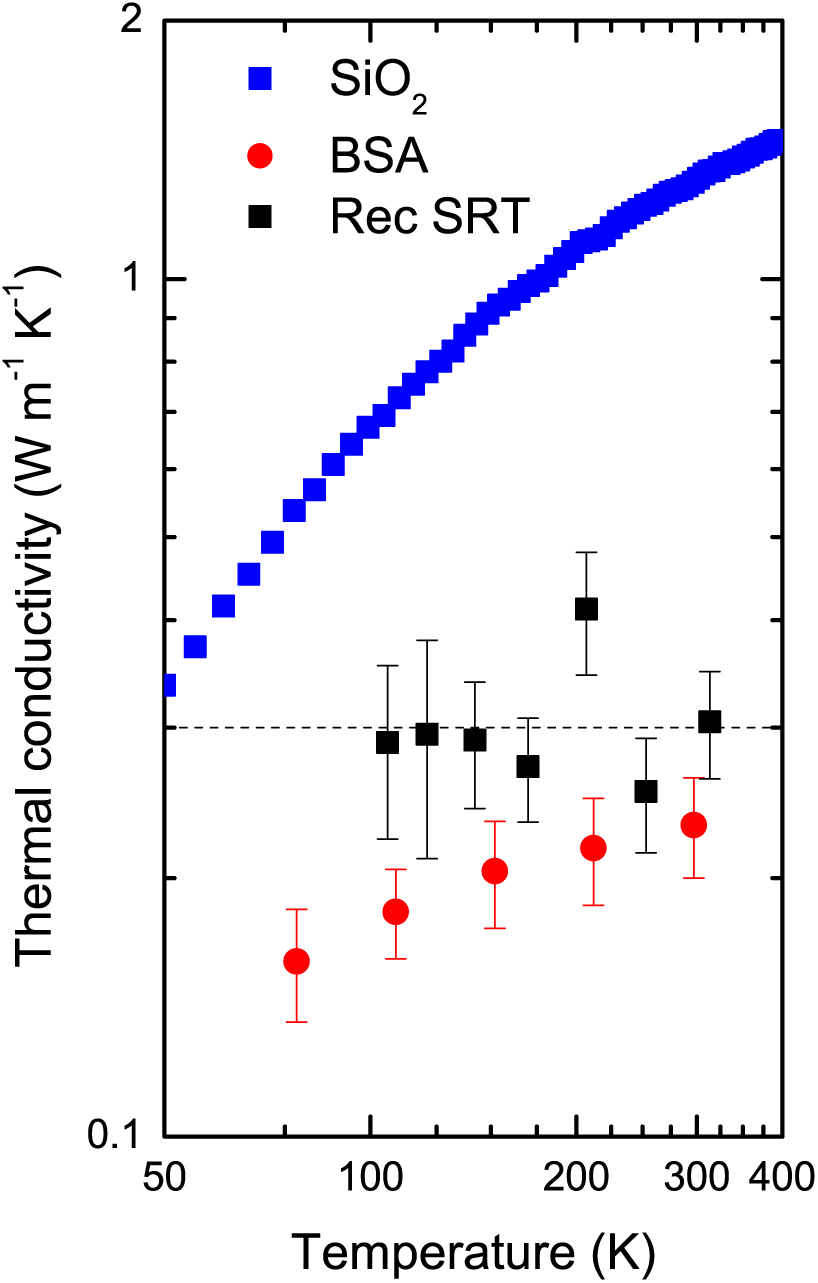
Thermal conductivity data of Silica (blue), crosslinked BSA (red) and recombinant SRT protein (black) are shown. Silica and crosslinked BSA are plotted for comparison and obtained from Hopkins et al. with permission. (B. M. Foley et al., J. Phys. Chem. Lett., 2014)

**Figure S6.**
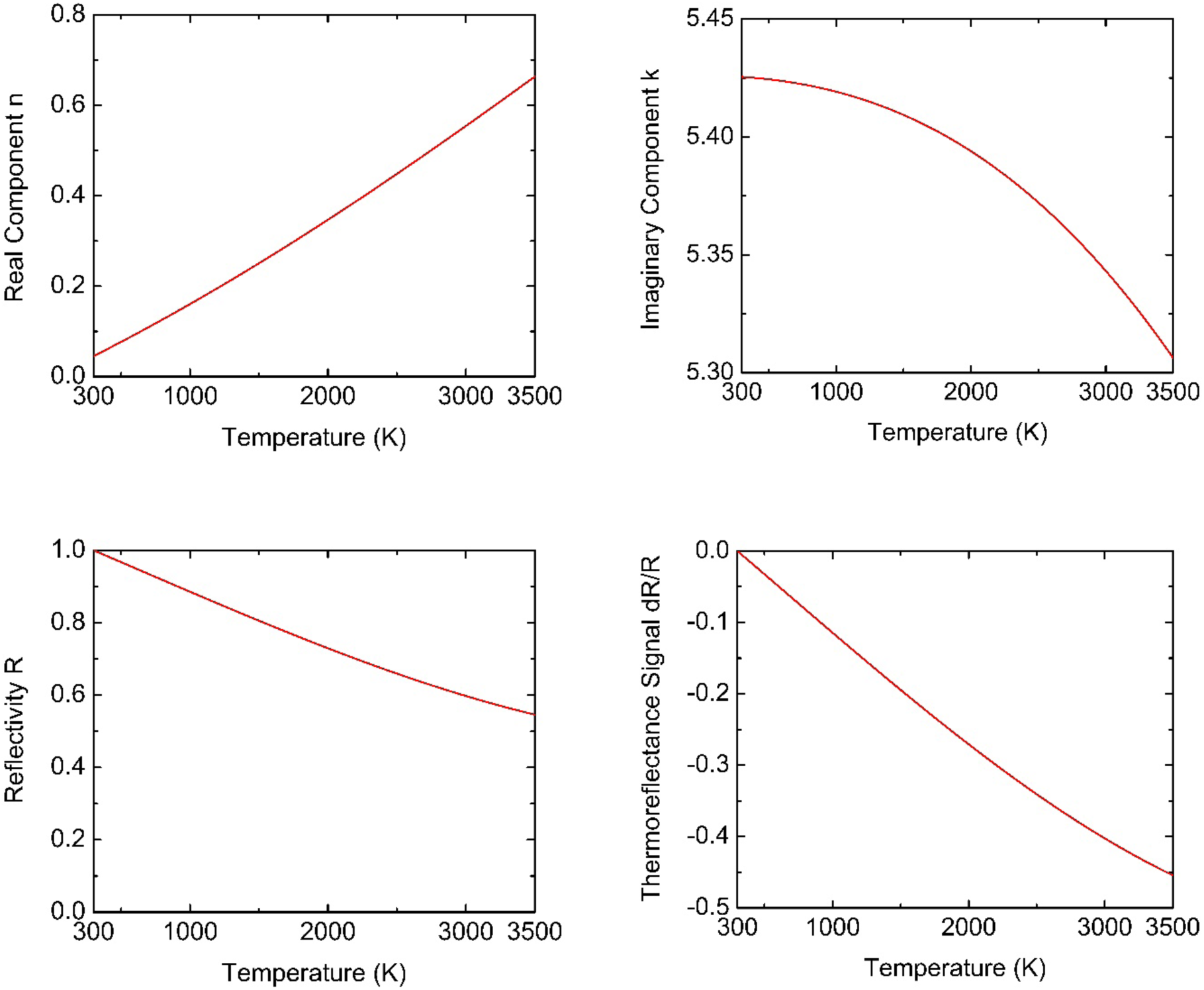
Calculated optical parameters of Au as a function of temperature as derived from the Drude model for the free electron complex dielectric function. (Top left) The real component of the refractive index, (top right) the imaginary component of the refractive index, (bottom left) the reflectivity, and (bottom left) the normalized change in reflectivity.

**Figure S7.**
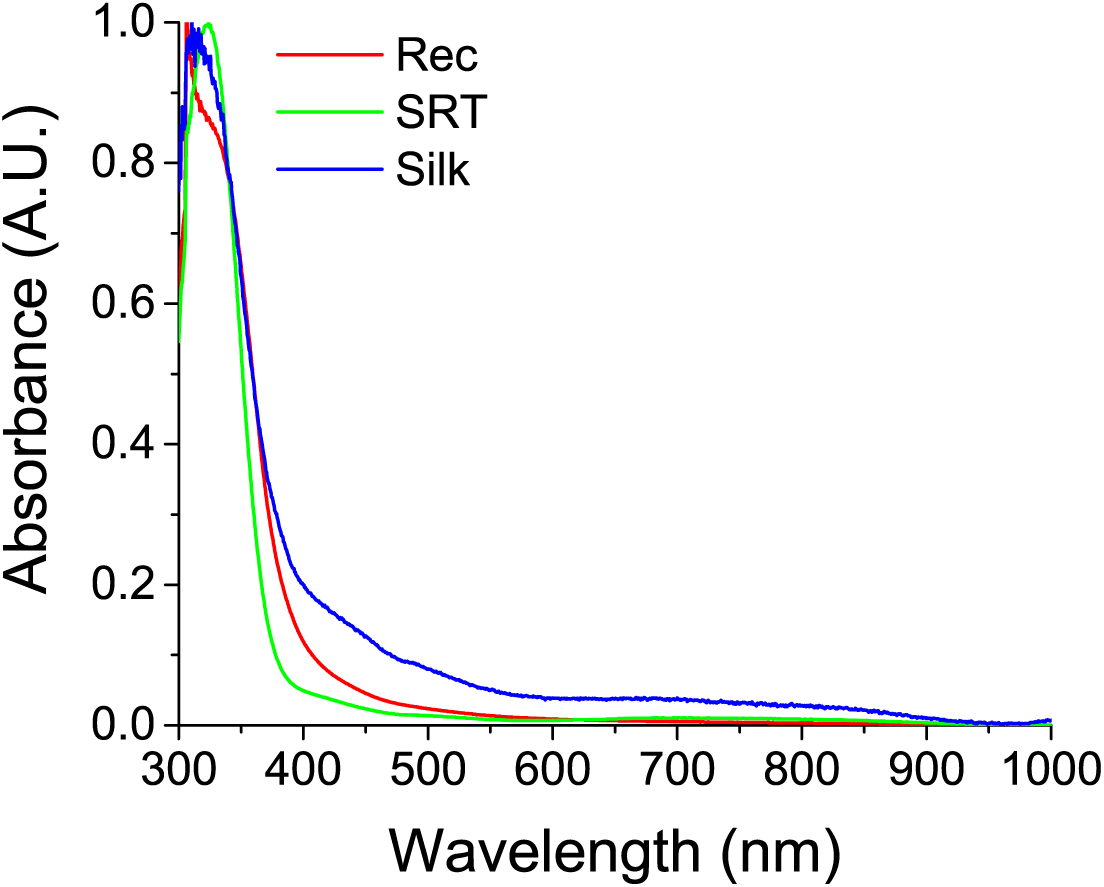
Optical absorbance of three proteins studied in this manuscript. High absorbance in 280-360nm regions is observed (as expected), whereas absorbance is low in 800 nm regions. We also note that typical laser stability of protein films are 1mW/mm^2^.

